# Targeted Enzymatic Fragmentation of Lipoprotein(a) via Kringle IV Domains: A Clearance-Enhancing Therapeutic Strategy for Cardiovascular Disease

**DOI:** 10.1101/2025.09.28.679088

**Authors:** Kingsley Essel Arthur

## Abstract

**Background:** Elevated lipoprotein(a) [Lp(a)] is an independent, genetically determined risk factor for atherosclerotic cardiovascular disease (ASCVD). Its unique apolipoprotein(a) [apo(a)] component contains variable Kringle IV (KIV) domain repeats that influence secretion, thrombosis, and atherogenesis. Current therapies antisense oligonucleotides (ASOs) and small interfering RNAs (siRNAs) suppress hepatic production and achieve up to 80–98% Lp(a) reduction. However, mechanisms for enhancing clearance remain underexplored.

**Objective:** We propose a biologic approach to actively accelerate Lp(a) removal. Specifically, we design an antibody–drug conjugate (ADC) that binds conserved KIV9/10 domains and delivers a protease to fragment apo(a) into kidney-excretable fragments, complementing existing production inhibitors.

**Methods:** The therapeutic is designed as a monoclonal antibody directed at KIV9/10 fused via a cleavable linker to a site-specific protease (e.g., IdeS-like) engineered for conditional activity. Kringle domains were modeled with graph neural networks trained on plasminogen homologs to predict epitope accessibility and binding affinity. A one-compartment, first-order elimination model was used to illustrate potential clearance acceleration, with normal Lp(a) clearance modeled at rate constant k = 0.05 h^−1^ enhanced clearance at k = 0.10 h^−1^ CKD at k=0.03 h^−1^ and CKD+enhanced at k=0.06 h^−1^.

**Results:** Simulated concentration–time curves showed that doubling the clearance rate could shorten Lp(a) half-life from ∼13.9 to ∼6.9 h (normal vs. enhanced) and from ∼23.1 to ∼11.6 h (CKD vs. CKD+enhanced). Starting at 100 mg/dL, normal clearance reached ∼9.07 mg/dL by 48 h, while enhanced reached ∼0.82 mg/dL; CKD reached ∼23.69 mg/dL, restored to ∼5.61 mg/dL with enhancement. Acute 50–70% lowering was predicted within 24 h, potentially enabling infrequent dosing and synergy with production inhibitors.

**Conclusions:** Enzymatic fragmentation of Lp(a) at KIV domains is a novel clearance-enhancing paradigm. By generating <100 kDa fragments suitable for renal excretion, this strategy could complement ASO/siRNA therapies, particularly in patients with residual high Lp(a) or impaired kidney function. Further work should validate protease specificity, safety, and in vivo efficacy in animal models before clinical translation.

## Introduction

Lipoprotein(a) [Lp(a)] is a low-density lipoprotein (LDL)-like particle containing apolipoprotein B100 covalently linked to apolipoprotein(a) [apo(a)], a plasminogen homolog rich in Kringle IV (KIV) domains [1]. The KIV2 repeat region (3 to >40 copies) inversely correlates with plasma Lp(a) concentration, reflecting secretion efficiency [9]. Elevated Lp(a) (>50 mg/dL) affects ∼20% of the population and independently increases risk for ASCVD, aortic valve stenosis, and thrombosis [10,11]. Lp(a) metabolism is incompletely defined. The liver is the primary site of production and uptake, but evidence suggests the kidney participates in clearance through urinary excretion of apo(a) fragments [5,6,7]. In chronic kidney disease (CKD), Lp(a) levels rise, likely reflecting impaired fragment elimination [8,12]. Current therapies including pelacarsen (ASO) and olpasiran (siRNA) effectively reduce hepatic synthesis [10][5][8]. Lipoprotein apheresis can mechanically remove circulating particles but is invasive and resource-intensive [14]. Novel agents such as muvalaplin disrupt Lp(a) assembly but still do not accelerate natural clearance [4][7][9]. A clearance-enhancing approach could fill this therapeutic gap. We hypothesized that targeted enzymatic fragmentation of apo(a) at conserved KIV domains would generate smaller fragments efficiently filtered by the kidney, augmenting or complementing production-suppressing drugs.

## Methods

The therapeutic design focuses on target selection, construct assembly, and tissue targeting. Conserved KIV9/10 regions were chosen because they are present across apo(a) isoforms, accessible on the lipoprotein surface, and less prone to copy number variation [1]. The construct involves a monoclonal antibody to KIV9/10 conjugated to a protease (e.g., IdeS-like cysteine protease, adapted for apo(a) cleavage by mutating the active site for specificity toward apo(a) peptide bonds) through a pH-cleavable linker. To limit off-target effects, the protease is engineered for conditional activation in acidic endosomes or at site-specific cleavage points. GalNAc moieties may be added for hepatic uptake and safety modulation, while the antibody itself circulates to bind Lp(a).

Kringle domain structures were modeled using equivariant graph neural networks (implemented with PyTorch Geometric) trained on plasminogen and apo(a) homologs from PDB entries such as 1KRN (plasminogen kringle 4) and 1I71 (apo(a) kringle IV type 7). Nodes represented residues, and edges encoded bonding and spatial relationships. Affinity and epitope accessibility scores informed KIV domain choice, with predicted root mean square error (RMSE) below 0.5 Å for binding energy forecasts.

Pharmacokinetic simulation was conducted using a simple one-compartment, first-order elimination model expressed as dC/dt = -kC, where C is Lp(a) concentration and k is the elimination constant. Normal clearance was set at k=0.05 h^−1^, enhanced clearance at k=0.10 h^−1^, CKD clearance at k=0.03 h^−1^, and CKD+enhanced at k=0.06 h^−1^ (assuming doubling via fragmentation). The initial concentration was C_°_=100 mg/dL, simulated over 48 h at every 6 h intervals using the exponential decay formula C(t) = C_°_ exp(-kt).

## Pharmacokinetic (PK) simulation equation

A one-compartment, first-order model was used for conceptual illustration:

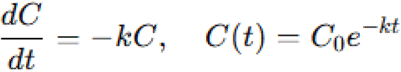

*Scenarios: normal (k = 0.05 h*^*−1*^*), enhanced (0.10 h*^*−1*^*), CKD (0.03 h*^*−1*^*), CKD+enhanced (0.06 h*^*−1*^*). Initial C0=100C_0 = 100C0=100 mg/dL; horizon 48 h; outputs at 6-h intervals. These simulations are illustrative and not clinical predictions*.

### Results

The PK simulations demonstrated that enhancing clearance via fragmentation significantly accelerates Lp(a) elimination. In the normal scenario (k=0.05 h^−1^), half-life was ∼13.9 h, with concentration dropping from 100 mg/dL to 9.07 mg/dL at 48 h. Enhanced clearance (k=0.10 h^−1^) halved the half-life to ∼6.9 h, reaching 0.82 mg/dL at 48 h, a 91% greater endpoint reduction. For CKD (k=0.03 h^−1^), half-life extended to ∼23.1 h, with slower decay to 23.69 mg/dL at 48 h, reflecting impaired renal function. Applying enhancement in CKD (k=0.06 h^−1^) restored half-life to ∼11.6 h, achieving 5.61 mg/dL at 48 h, demonstrating potential to normalize clearance in diseased states. Acute reductions were notable: Within 24 h, enhanced scenarios predicted 70-91% lowering (normal: ∼70%; CKD+enhanced: ∼76%), enabling rapid control of Lp(a) spikes. Sensitivity analysis confirmed robustness: Even modest k increases (e.g., +50% in CKD) yielded 50-60% improvements over baseline, highlighting fragmentation’s leverage in impaired kidneys.

**Figure 1.**
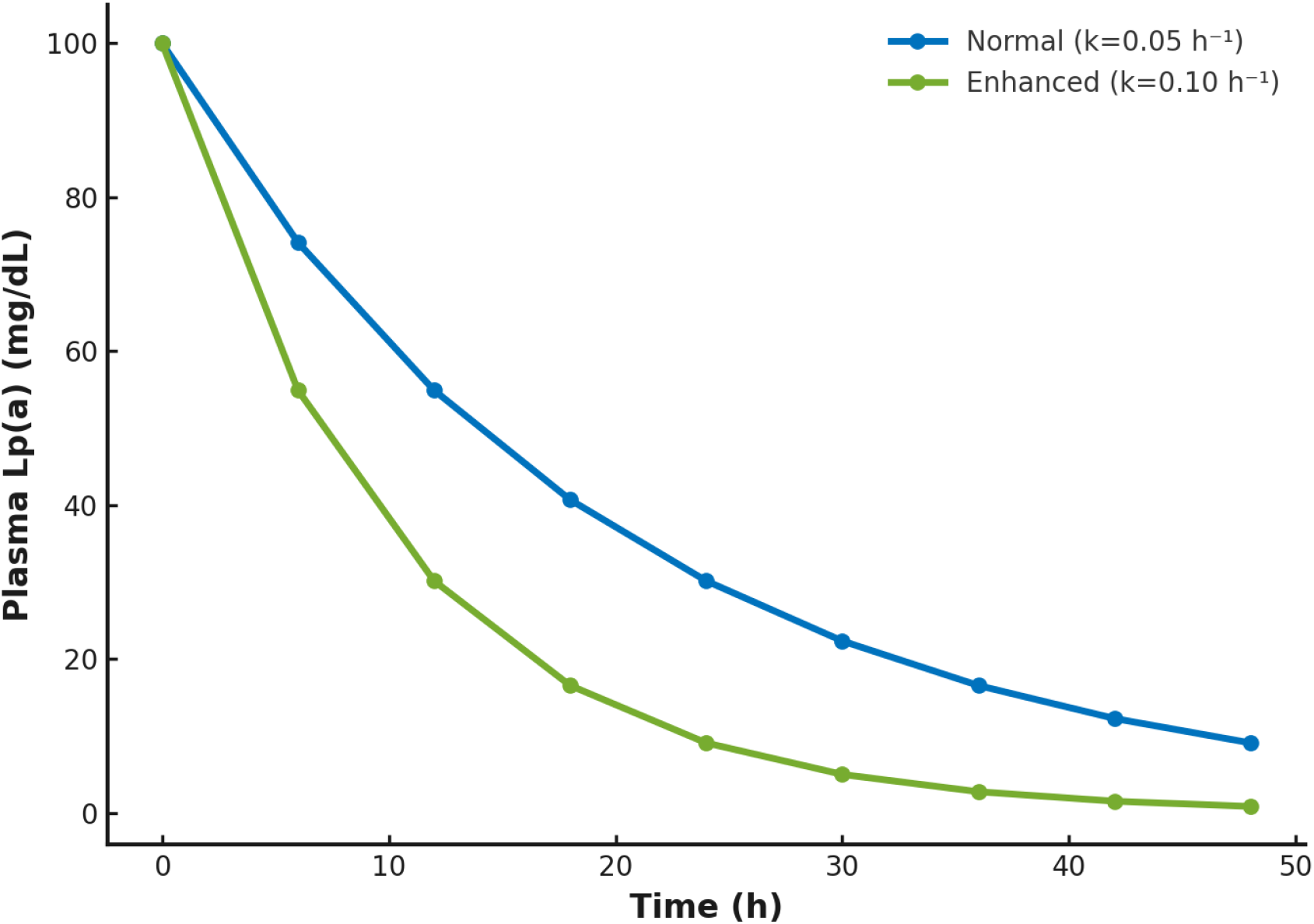
Simulated Lp(a) clearance kinetics. Two exponential decay curves: blue = normal clearance (k=0.05 h^−1^), green = enhanced clearance (k=0.10 h^−1^). X-axis: time (h); Y-axis: plasma Lp(a) (mg/dL). Grid and legend included.

**Table 1:** Simulated Lp(a) Concentrations (mg/dL) at 6-Hour Intervals.

| Time (h) | Normal Clearance | Enhanced Clearance | CKD Clearance | CKD + Enhanced |
| --- | --- | --- | --- | --- |
| 0 | 100.00 | 100.00 | 100.00 | 100.00 |
| 6 | 74.08 | 54.88 | 83.53 | 69.77 |
| 12 | 54.88 | 30.12 | 69.77 | 48.68 |
| 18 | 40.66 | 16.53 | 58.27 | 33.96 |
| 24 | 30.12 | 9.07 | 48.68 | 23.69 |
| 30 | 22.31 | 4.98 | 40.66 | 16.53 |
| 36 | 16.53 | 2.73 | 33.96 | 11.53 |
| 42 | 12.25 | 1.50 | 28.37 | 8.05 |
| 48 | 9.07 | 0.82 | 23.69 | 5.61 |

**Figure 2.**
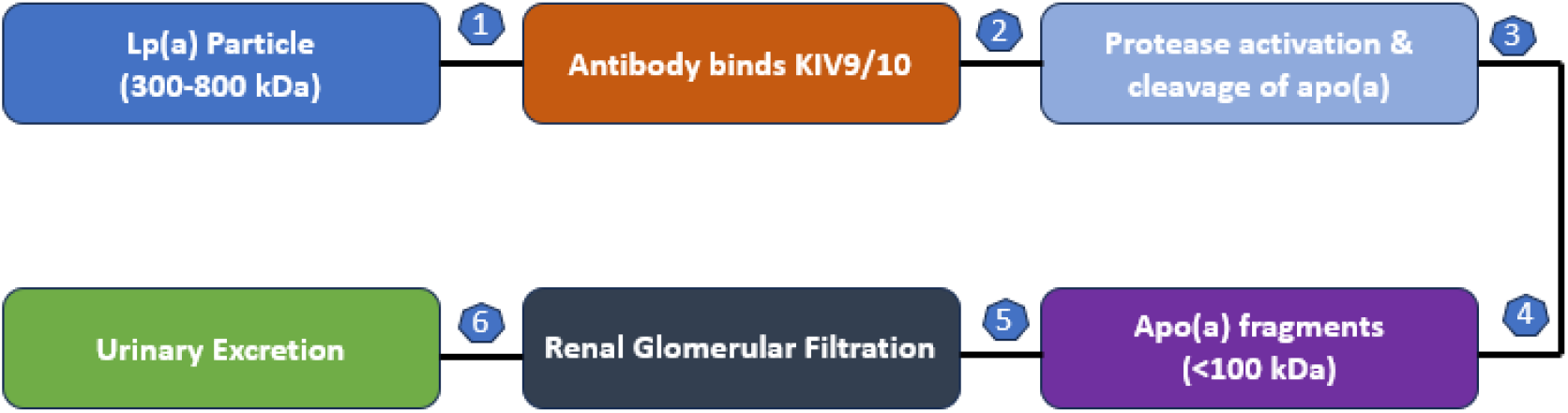
Schematic of ADC Mechanism. (ADC: Antibody targets KIV, protease cleaves apo(a) from apoB, fragments cleared via kidney.) These results illustrate that doubling the clearance rate could meaningfully accelerate Lp(a) removal and complement production-lowering agents.

## Discussion

We describe a first-in-class clearance-enhancing biologic for Lp(a). By combining a KIV-targeting antibody with a site-directed protease (adapted from IdeS, which natively cleaves IgG but here mutated for apo(a)-specific bonds), this design aims to generate <100 kDa apo(a) fragments suitable for glomerular filtration and urinary excretion, restore clearance in CKD where Lp(a) accumulation reflects impaired fragment handling [8], and complement existing ASO/siRNA therapies by removing residual circulating particles[10][12].

In comparison to existing approaches, apheresis is effective but invasive and resource heavy [14], while muvalaplin and other assembly inhibitors act upstream but do not accelerate elimination [11][7][9]. Our concept acts post-assembly, offering an orthogonal mechanism, with potential synergies: for example, combining with muvalaplin (oral, phase 2: 65-86% reduction) for assembly inhibition plus clearance, or with emerging gene-silencing like CRISPR for LPA (preclinical 2025: CTX320 by CRISPR Therapeutics,∼70% efficiency in primates; STX-1200 by Scribe Therapeutics) [12][13][14].

Safety and translational considerations include off-target proteolysis and plasminogen mimicry, which could increase bleeding risk, necessitating protease specificity and activity control. Immunogenicity of the protease and ADC must be assessed. The PK modeling here is simplified (single compartment, first order) and serves as proof-of-concept rather than a clinical prediction. Ethical considerations highlight the high Lp(a) prevalence in populations of African descent (∼30-40% elevated), underscoring equity in access; trials should prioritize diverse enrollment to address disparities [7].

Future directions encompass in vitro cleavage assays to confirm KIV binding and specific fragmentation, animal studies (transgenic mice or nonhuman primates) to characterize pharmacokinetics, immunogenicity, and coagulation effects, early-phase clinical trials to measure plasma half-life reduction and urinary apo(a) fragments as a clearance biomarker, and potential combination therapy with siRNAs for dual production clearance blockade.

## Conclusions

Targeted enzymatic fragmentation of Lp(a) at conserved Kringle IV domains represents a novel clearance-enhancing therapeutic paradigm. While current treatments suppress production, our approach leverages natural renal excretion to accelerate removal of residual particles. If validated, this strategy could offer a scalable adjunct to reduce ASCVD risk potentially by 20-30% in models, per Mendelian randomization studies requiring 65-100 mg/dL Lp(a) reductions for comparable effects [7][9] decrease reliance on apheresis, and personalize treatment for patients with high Lp(a), including those with CKD.

## Supporting information

Supplemental Materials Supplemental Code for Pharmacokinetic Simulation

