## Supplemental Materials Supplemental Code for Pharmacokinetic Simulation for "Targeted Enzymatic Fragmentation of Lipoprotein(a) via Kringle IV Domains: A Clearance-Enhancing Therapeutic Strategy for Cardiovascular Disease"

**Supplemental Code for Pharmacokinetic Simulation**

To ensure reproducibility, the following Python code was used to generate the pharmacokinetic simulations and table data. It requires NumPy (for calculations) and Matplotlib (for plotting) and can be run in a standard Python environment. Error handling is added for invalid inputs (e.g., negative k or C0), and export options include saving the table to CSV and the plot to PNG.

---

```
import numpy as np

import matplotlib.pyplot as plt

import csv


# Parameters with error handling

try:

    C0 = 100  # initial concentration (mg/dL)

    if C0 <= 0:

        raise ValueError("Initial concentration C0 must be positive.")


    t_table = np.arange(0, 49, 6)  # 6-hour intervals up to 48 h

    k_normal = 0.05

    k_enhanced = 0.10

    k_ckd = 0.03

    k_ckd_enhanced = 0.06
```

```
    # Validate k values
    ks = [k_normal, k_enhanced, k_ckd, k_ckd_enhanced]
    if any(k <= 0 for k in ks):
        raise ValueError("All rate constants k must be positive.")

    # Calculate concentrations
    C_normal = C0 * np.exp(-k_normal * t_table)
    C_enhanced = C0 * np.exp(-k_enhanced * t_table)
    C_ckd = C0 * np.exp(-k_ckd * t_table)
    C_ckd_enhanced = C0 * np.exp(-k_ckd_enhanced * t_table)

    print("Time (h):", t_table)
    print("Normal:", np.round(C_normal, 2))
    print("Enhanced:", np.round(C_enhanced, 2))
    print("CKD:", np.round(C_ckd, 2))
    print("CKD + Enhanced:", np.round(C_ckd_enhanced, 2))

    # Export table to CSV
    with open('lp_a_concentrations.csv', 'w', newline='') as csvfile:
        writer = csv.writer(csvfile)
        writer.writerow(['Time (h)', 'Normal Clearance', 'Enhanced Clearance', 'CKD Clearance', 'CKD + Enhanced'])
```

```
        for row in zip(t_table, np.round(C_normal, 2), np.round(C_enhanced, 2), np.round(C_ckd, 2), np.round(C_ckd_enhanced, 2)):
            writer.writerow(row)
    print("Table exported to 'lp_a_concentrations.csv'")


    # Plot curves for Figure 1
    t = np.linspace(0, 48, 100)
    plt.figure(figsize=(10, 6))
    plt.plot(t, C0*np.exp(-k_normal*t), label='Normal (k=0.05 h⁻¹)', color='blue')
    plt.plot(t, C0*np.exp(-k_enhanced*t), label='Enhanced (k=0.10 h⁻¹)', color='green')
    plt.plot(t, C0*np.exp(-k_ckd*t), label='CKD (k=0.03 h⁻¹)', color='red', linestyle='--')
    plt.plot(t, C0*np.exp(-k_ckd_enhanced*t), label='CKD+Enhanced (k=0.06 h⁻¹)', color='orange', linestyle='--')
    plt.xlabel('Time (h)')
    plt.ylabel('Plasma Lp(a) (mg/dL)')
    plt.title('Simulated Lp(a) Clearance Kinetics')
    plt.legend()
    plt.grid(True)
    plt.savefig('lp_a_kinetics.png')
    plt.show()
    print("Plot exported to 'lp_a_kinetics.png'")

except ValueError as e:
```

```
    print(f"Error: {e}")
except Exception as e:
    print(f"Unexpected error: {e}")
```

---

This code produces the exact values in Table 1, exports the table as CSV for data sharing, and saves the plot as PNG for easy inclusion in documents. All simulations are deterministic and reproducible with the given parameters. If errors occur (e.g., invalid parameters), it handles them gracefully without crashing.
